# Status and Genetic Diversity of *Hipposideros pratti* in the Northernmost Part of Its Distribution

**DOI:** 10.1101/2023.07.15.549176

**Authors:** Yanmei Wang, Luwen Shang, Jinhe Wang, Liming Zhao, YanZhen Bu

## Abstract

*Hipposideros pratti* is species of bat distributed in caves from Southeast Asia to the Qinling-Funiu Mountain area in China. To understand the dynamic changes in its distribution and evaluate the current health status of the species, we reviewed 48 caves in the northernmost area of the *H. pratti* distribution and conducted the first analysis of genetic diversity for the species in the region. *H. pratti* was only detected in three caves, the four previously distributed caves and the remaining 41 previously undistributed caves were not observed. An analysis based on microsatellite markers revealed a few key points. 1) The average number of observed alleles (*N*_*a*_) of *H. pratti* in the region was 3.94, and the average observed heterozygosity (*H*_*o*_) was 0.5293. All three populations were in Hardy–Weinberg equilibrium. 2) Intra-population was the predominant genetic variation of the *H. pratti* population. 3) *H. pratti* in the region experienced a bottleneck effect. We found that the three caves where *H. pratti* are currently distributed face varying degrees of human interference and the populations are threatened. Management strategies, including appropriate countermeasures to reduce human interference in caves, are urgently needed.

## 1 INTRODUCTION

*Hipposideros pratti* belongs to the family Hipposideridae of Chiroptera and is widely distributed in tropical and subtropical regions of Asia. The Qinling-Funiu Mountain area is the northernmost boundary of its distribution (Robinson *et al*.,2003); this area is the boundary between the Palaearctic and Oriental realms (Chen, 2004). Studies of *H. pratti* have mainly focused on gastrointestinal bacteria (Yuan *et al*., 2017), echolocation (Yang, 2018), embryonic development (Wang *et al*., 2010), and functional gene evolution (Gang *et al*., 2006); however, detailed analyses of genetic diversity in this species are lacking.

Genetic diversity provides a theoretical basis for judging the current status of species and the degree of genetic diversity is related to evolutionary trends (Cheng *et al*., 2015). Studies of genetic diversity in *H. pratti* are necessary for the formulation of scientific and effective protection strategies and management methods for the species. Therefore, we investigated the region in the northernmost segment of its distribution and evaluated genetic diversity. From 2011 to 2013, we conducted a systematic survey of the area and found seven caves with *H. pratti*. Further investigations of this area will be performed to monitor the population dynamics and health status of *H. pratti*.

## 2 MATERIALS AND METHODS

### 2.1 Overview of the survey area

Funiu Mountain (E 110°30′−113°05′, N 32°45′−34°00′) is mainly located in Nanyang City, Henan Province and is located at the boundary between subtropical and warm temperate zones in China (Zhang and Mu, 2021). It belongs to the national nature reserve and is the northernmost tip of the distribution of *H. pratti*. The terrain is roughly tilted from northwest to southeast. Due to its special geographical location and complex and diverse environment, it is rich in natural resources (He *et al*., 2021). There are many caves in Funiu Mountain as well as some tunnels and abandoned mines, which have become the main habitat of cave bats (Wu *et al*., 2018). When we conducted our survey in 2011–2013, 48 caves were investigated, of which 7 had *H. pratti* (Bu *et al*., 2014). In this study, 48 caves will be investigated again.

### 2.2 Specimen collection

A survey was carried out in caves in the Funiu Mountain area. If *H. pratti* was detected, the wing membrane was sampled with a punch, temporarily stored in 95% alcohol, and stored in a refrigerator for subsequent experiments. After sampling, the individual was released in situ.

### 2.3 Extraction of genomic DNA

The DNA Extraction Kit produced by Tiangen was used to extract DNA from the wing membrane samples of *H. pratti*. The concentration and purity of DNA were detected and recorded by ultraviolet spectrophotometry and agarose gel electrophoresis. Then, samples were diluted to 100 ng/μL and stored at -20°C.

### 2.4 Selection of microsatellite markers and PCR amplification

Six pairs of microsatellite primers were used to evaluate genetic diversity in *H. pratti* (Tables 1). The PCR amplification system and conditions were optimized through multiple experiments, and six pairs of microsatellite primers were used for PCR amplification for each individual in each population. The total PCR amplification system was 25 μL, containing DNA template (1.3 μL), 2× Taq Plus PCR Mix (12.5 μL), Primer F (0.65 μL), and Primer R (0.65 μL), ddH_2_O (9.9 μL). The optimal PCR conditions were as follows: initial denaturation at 95 °C for 5 min, 34 cycles of denaturation at 95 °C for 30 s, annealing for 30 s, and extension at 72 °C for 1 min, and extension at 72 °C for 10 min.

**Table 1.** *H. pratti* sample collection sites.

| Collection sites | Number | Latitude | Longitude | Altitude ( m ) |
| --- | --- | --- | --- | --- |
| XX | 3 | 33°18'N | 111°25'W | 415 |
| TX | 9 | 33°20'N | 111°52'W | 289 |
| XR | 4 | 33°34'N | 112°06'W | 349 |

### 2.5 Genotyping and target fragment acquisition

Six pairs of microsatellite primers were fluorescently labeled at the 5’ end, and fluorescent groups were selected according to the size of the target fragment to synthesize fluorescent primers (Tables 1). The fluorescent products were detected by agarose gel electrophoresis. The three fluorescent signals for each individual were mixed in proportion (FAM: HEX: TAMRA = 1:2:2) and sent to Jinweizhi Biological Company for capillary electrophoresis (ABI3700 sequencing instrument). Based on the genotyping results, allele information was used to identify whether the target fragment is homozygous or heterozygous.

### 2.6 Population genetic analyses

PopGene 1.32, FSTAT 2.9.3, and Genepop 4.7.5 were used to analyze the genetic diversity in the *H. pratti* population. The observed number of alleles (*N*_*a*_), effective number of alleles (*N*_*e*_), Shannon information index (*I*), observed heterozygosity (*H*_*O*_), and expected heterozygosity (*H*_*e*_) were calculated using PopGene 1.32. Invalid allele frequency, Hardy-Weinberg equilibrium and linkage disequilibrium were calculated using Genepop 4.7.5. The allelic richness (AR) and inbreeding coefficient (*F*_IS_) were calculated using FSTAT 2.9.3. Arlequin 3.5 and FSTAT 2.9.3 were used to analyze genetic variation. Arlequin 3.5 was used to calculate the genetic differentiation coefficient (*F*_ST_) and gene flow (*Nm*) between populations. Analysis of molecular variance (AMOVA) was used to analyze the source of population variation. FSTAT 2.9.3 was used to calculate Nei’s gene differentiation coefficient. Bottleneck 1.2.2 was used to calculate the gene frequency distribution model of *H. pratti* with a repetition of 1000 to detect bottleneck events under the two-phase model of mutation (TPM).

## 3 RESULTS

### 3.1 Survey results

From May to October 2019, the distribution of *H. pratti* in caves in the Funiu Mountain area was investigated again. Among 48 caves, *H. pratti* was found only in Xixia Cave (XX), Tianxin Cave (TX), and Xianren Cave (XR) (Fig. 1). In four caves with bat records between 2011 and 2013 (Banchang, Shangji, Junmahe, and Sunmen), *H. pratti* was not found in this survey. Additionally, 41 caves with no previous records of *H. pratti* still lacked the species. A total of 16 bats were captured (Table 1).

**Fig. 1.**
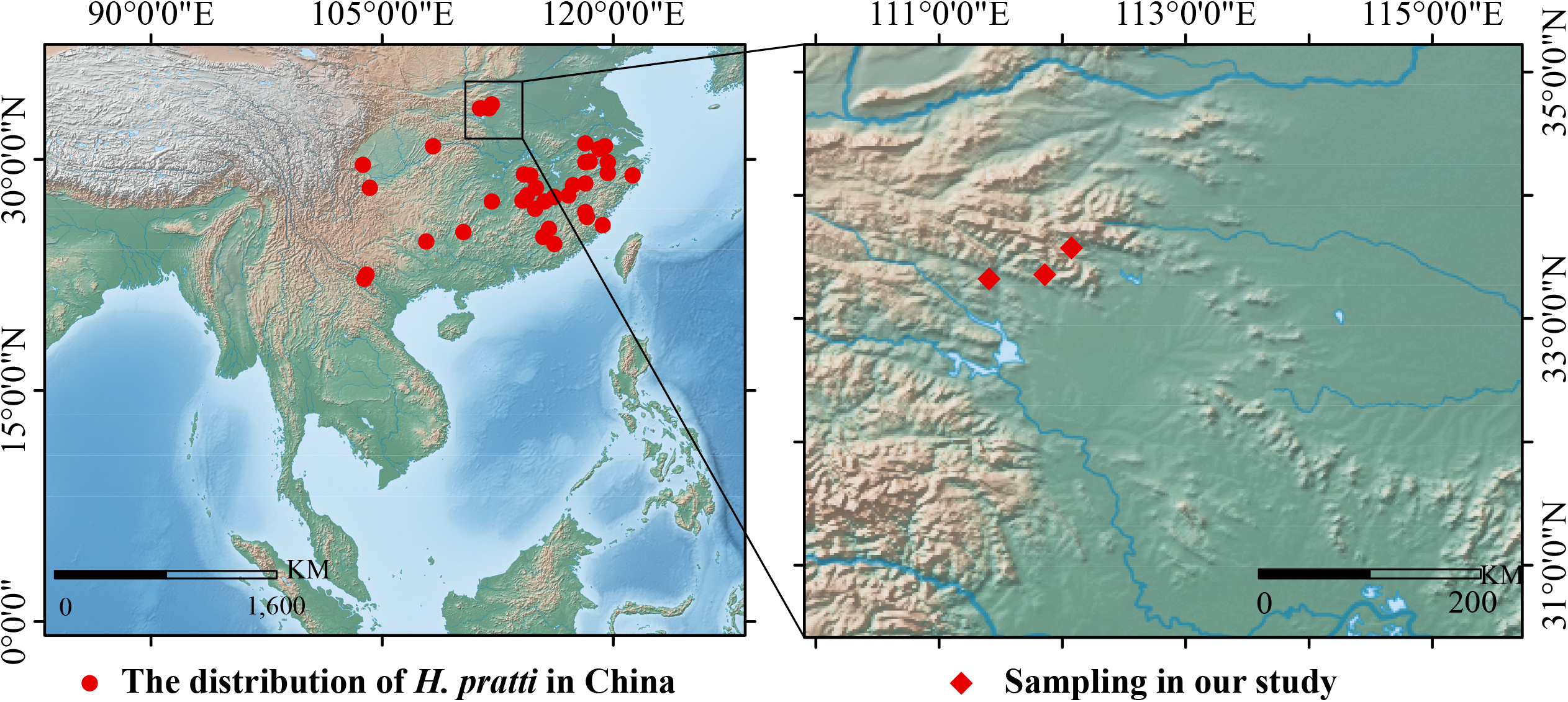
Northernmost tip of the *H. pratti* distribution. Note: The circle represents the distribution of *H. pratti*. Undetected twice: In both surveys, *H. pratti* was not detected. For the first time: From 2011 to 2013, caves within the *H. pratti* distribution were observed for the first time. Detected twice: From 2011 to 2013 and 2019, caves were within the distribution of *H. pratti*.

### 3.2 Population genetic diversity

A total of 48 alleles were amplified at 6 microsatellite loci in 16 individuals from three *H. pratti* populations, with 3–18 alleles per locus (Tables 2). The overall average observed number of alleles (*N*_*a*_) in the three populations was 3.94, and the overall observed heterozygosity was close to the expected heterozygosity. In the three populations, the inbreeding coefficient (*F*_IS_) for the Xixia population (XX) was less than 0, and the population had excess heterozygotes. The inbreeding coefficients for the other populations were greater than 0, with had heterozygote deficiencies (Table 2). A linkage disequilibrium test showed that the selected microsatellite loci were inherited independently. Among the six loci, the Xixia population (XX) had only one allele at the P5 and PT loci; accordingly, we could not evaluate deviations from the Hardy–Weinberg equilibrium. At the other loci in each population, deviations from Hardy–Weinberg equilibrium were not detected (Tables 3). In view of this, invalid alleles for each population at each locus were detected. For loci P5 and PT, the Xixia population (XX) had only one allele and therefore no invalid alleles were detected (Tables 4).

**Table 2.**
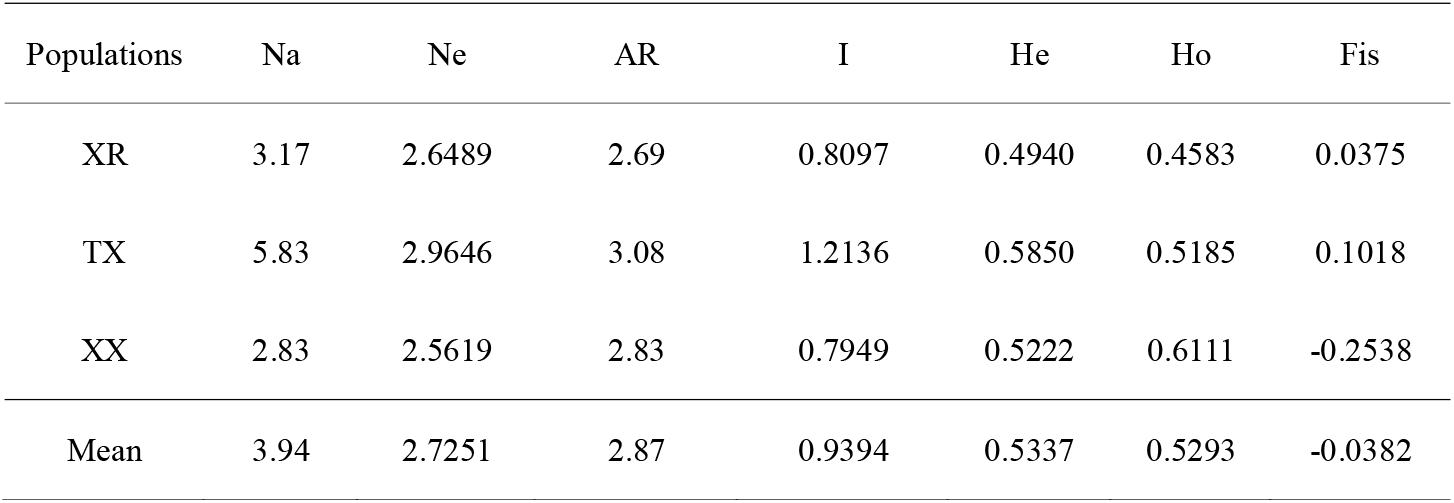
Genetic diversity in *H. pratti* populations.

| Populations | Na | Ne | AR | I | He | Ho | Fis |
| --- | --- | --- | --- | --- | --- | --- | --- |
| XR | 3.17 | 2.6489 | 2.69 | 0.8097 | 0.4940 | 0.4583 | 0.0375 |
| TX | 5.83 | 2.9646 | 3.08 | 1.2136 | 0.5850 | 0.5185 | 0.1018 |
| XX | 2.83 | 2.5619 | 2.83 | 0.7949 | 0.5222 | 0.6111 | -0.2538 |
| Mean | 3.94 | 2.7251 | 2.87 | 0.9394 | 0.5337 | 0.5293 | -0.0382 |

**Table 3.** Analysis of molecular variance (AMOVA) based on microsatellite data.

| Source of variation | Sum of squares | Variance components | Percentage variation |
| --- | --- | --- | --- |
| Among populations | 4.083 | 0.02872 | 1.69309 |
| Among individuals<br>within populations | 23.042 | 0.10497 | 6.18847 |
| Within individuals | 25.000 | 1.56250 | 92.11844 |

### 3.3 Genetic variation in *H. pratti* populations

The coefficients of genetic differentiation (*F*_ST_) between the two populations were 0.00514–0.04485. The Xixia (XX) and Xianren (XR) populations had the highest degree of differentiation and genetic variation, with a maximum *F*_ST_ of 0.04485. At the same time, divergence between Xixia (XX) and Tianxin (TX) populations was the lowest, and *F*_ST_ was 0.00514 (Tables 5).

Gene flow (*Nm*) estimates between the two populations exceeded 1. The Nm in the Xixia (XX) and Xianren (XR) populations was the lowest (5.3241). There was frequent gene exchange between the Xixia (XX) and Tianxin (TX) populations, the Nm estimates reaching 48.3881 (Tables 5).

In the three populations, the total genetic diversity (*H*_T_) was 0.546, the average genetic diversity (*H*_S_) within populations was 0.536, and the average genetic diversity (*D*_ST_) between populations was 0.01. The genetic diversity was dominated by variation within the population, accounting for 98.17% of the total variation. The coefficient of genetic differentiation (*G*_ST_) for the three populations at six polymorphic loci ranged from -0.062 to 0.075, with an average of 0.018 (Tables 6).

AMOVA revealed that inter-regional genetic variation accounted for 1.69309% of the total genetic variation, and the genetic variation within populations accounted for 92.11844% (Table 3).

### 3.4 Bottleneck effects

The bottleneck effect in the *H. pratti* population in Funiu Mountain was analyzed. Wilcoxon’s test suggested that there was a significant bottleneck effect in the population (P = 0.01563) under the second-order mutation model (TPM).

## 4 DISCUSSION

### 4.1 Reduced range of distribution in the northernmost tip of the *H. pratti*

*H. pratti* has strong selectivity for habitats. It is a typical cave bat and is often found at the top of the cave near the exit (Wang *et al*., 2021). The species is widely distributed, according to a national survey, it is found in Jiangxi, Fujian, Guangdong, and other central and southern provinces (He, 2015). *H. pratti* has been widely distributed at the northernmost end of the past (Niu *et al*., 2010). In our 2011–2013 survey in the Funiu Mountain area, *H. pratti* was found in seven caves; however, only three remain. The three caves have been developed for tourism, resulting in a sharp decline in the number of bats. The negative impacts of modernization-related construction, such as urbanization construction, high-speed railway construction, and cave tourism development, may explain the decrease of the distribution of *H. pratti*.

### 4.2 Inbreeding in *H. pratti* populations

Observed heterozygosity (*Ho*) and expected heterozygosity (*He*) indicate the heterozygosity of alleles at a certain locus in the population, and higher values indicate richer genetic diversity (Zou *et al*., 2015). Values of 0.3 □ *H*_*e*_ □ 0.8 indicates that the population has some degree of genetic diversity, and *H*_*o*_ □ *H*_*e*_ indicates that inbreeding occurred within the population (Takezaki and Nei, 1996; Huang *et al*., 2005). The expected heterozygosity (*H*_*e*_) in the three populations was 0.4940–0.5850, and the observed heterozygosity (*H*_*o*_) was 0.4583–0.6111. Overall, the *H*_*E*_ values of the three populations in Funiu Mountain were generally greater than the *H*_*o*_ values, indicating that there was inbreeding.

In the three populations, the inbreeding coefficients (*F*_IS_) of Tianxin (TX) and Xianren (XR) were greater than 0, indicating that these populations had heterozygote deficiencies, consistent with inbreeding. Typically, inbreeding in population genetics has negative effects on long-term evolution. Inbreeding and genetic drift within populations can result in deviations from Hardy–Weinberg equilibrium (Hartl, 2020). Based on 18 Hardy-Weinberg equilibrium tests at six loci, the three populations conformed to the Hardy–Weinberg equilibrium, indicating that the gene frequencies in the *H. pratti* populations in Funiu Mountain were relatively stable and a balance was maintained across generations.

### 4.3 Genetic differentiation in *H. pratti*

The genetic differentiation coefficient (*F*_ST_) reflects the genetic differentiation level of population (Wright, 1978). The coefficients of genetic differentiation (*F*_ST_) for *H. pratti* populations were 0.00514–0.04485, indicating that the degree of genetic differentiation among populations was low (Curnow and Wright, 1979). Gene flow (*Nm*) is the main factor limiting genetic differentiation among populations. *Nm* □ 1 indicates gene flow between populations (Wright, 1990), as observed between the three populations in this study. According to Nei ‘s gene differentiation and AMOVA, the genetic variation within the population was the largest determinant of overall variation in *H. pratti*.

### 4.4 *H. pratti* bottleneck

*H. pratti* populations in the northernmost part of the species distribution may have experienced a bottleneck event, resulting in a decline in genetic diversity in the area. However, the current environmental conditions still pose a great threat to the survival of *H. pratti*. Therefore, real-time monitoring is needed to evaluate the population size and dynamics and thereby to avoid habitat destruction caused by human disturbance conserve genetic diversity in *H. pratti* in Funiu Mountain, in the northernmost part of China. A greater awareness of the protection of this species is needed to minimize human disturbance near the only three caves in which it has been observed.

## 5 CONCLUSIONS

In summary, Caves with *H. pratti* in the northernmost area of its distribution decreased, and the habitat was seriously disturbed by human beings; Inbreeding was detected in populations of *H. pratti* in the northernmost part of its distribution; Both gene flow and genetic differentiation were detected among the three populations, and intra-population was the predominant genetic variation of the *H. pratti* population; *H. pratti* in the area experienced a significant bottleneck.

## SUPPORTING INFORMATION

**Additional file 1**: Additional Information of article. (DOC 115kb)

**Additional file 2**: All data analysed during this study. (XLS 20.5kb)

## ACKNOWLEDGMENTS

This project was supported by the Study on the Diversity of Chiroptera in Henan Province and the National Natural Science Foundation of China (No. U1704102).

## DATE ACCESSIBILITY STATEMENT

All data generated or analyzed during this study are included in this published article [and its supplementary information files].

## AUTHORS CONTRIBUTIONS

Yanmei Wang and YanZhen Bu designed the research; Luwen Shang and Jinhe Wang collected the field data; Liming Zhao analyzed the data and made the figures; Yanmei Wang and YanZhen Bu led the writing of the manuscript. All authors contributed intellectually to the discussion framework of the manuscript and gave final approval for publication.

